# Inhibition of epithelial Na^+^ transport: novel mechanism of *Ureaplasma*-driven lung disease

**DOI:** 10.1101/2024.02.14.580256

**Authors:** Kirsten Glaser, Carl-Bernd Rieger, Elisabeth Paluszkiewicz, Ulrich H. Thome, Mandy Laube

## Abstract

**Background:** Respiratory tract colonisation with *Ureaplasma* species has been associated with the development of acute and long-term pulmonary morbidity in preterm infants. Apart from inflammation, the underlying mechanisms of *Ureaplasma*-driven lung disease are mainly unknown. The present investigation is the first to examine the influence of acute *Ureaplasma* infection on critical mechanisms of alveolar fluid clearance in the immature lung.

**Methods:** Primary rat fetal distal lung epithelial (FDLE) cells were incubated with viable *Ureaplasma* in the absence or presence of the urease inhibitor flurofamide. Na^+^ transport and activity of the epithelial Na^+^ channel (ENaC) and the Na,K-ATPase were determined in Ussing chambers. Barrier integrity, metabolic activity, gene expression, and kinase signalling were also assessed.

**Results:** We found a 30-90% decrease of epithelial Na^+^ transport upon 24 hours of *Ureaplasma* infection resulting from significant inhibition of ENaC and Na,K-ATPase activities. Notably, *Ureaplasma* induced phosphorylation of Erk1/2 – a well-known inhibitor of ENaC activity. Moreover, *Ureaplasma-*driven NH_3_ production - and not the accompanying pH shift - inhibited the epithelial Na^+^ transport. Co-incubation with flurofamide entirely restored Na^+^ transport in *Ureaplasma*-infected FDLE cells.

**Conclusion:** Our data demonstrate that *Ureaplasma* infection significantly impairs epithelial Na^+^ transport and subsequent fluid clearance in fetal alveolar cells – most likely by Erk1/2 phosphorylation. We identified NH_3_ as the mediating virulence factor and were able to restore Na^+^ transport by inhibiting the *Ureaplasma*-specific urease. This is the first study to show a functional impairment of pulmonary epithelial cells upon *Ureaplasma* infection, revealing a potential mechanism of *Ureaplasma*-driven preterm lung disease.

**Take Home:** We report *Ureaplasma*-induced inhibition of epithelial Na^+^ transport as a potential mechanism of *Ureaplasma*-driven preterm lung disease. NH_3_ is identified as a virulence factor offering a potential therapeutic role for urease inhibitors in colonised infants.

## INTRODUCTION

*Ureaplasma* species (spp.) are mostly commensal bacteria of the adult urogenital tract. Yet, in pregnancy, *Ureaplasma (U.) parvum* and *U. urealyticum* have been associated with chorioamnionitis and preterm birth [1]. In preterm infants, epidemiological and experimental data indicate a strong correlation between *Ureaplasma* respiratory tract colonisation and adverse pulmonary short- and long-term outcomes, including the development of bronchopulmonary dysplasia [2, 3]. Colonisation rates increase with decreasing gestational age, leaving the most preterm infants at highest risk [3]. Animal studies and *in vitro* data confirm inflammatory responses and altered lung development in *Ureaplasma*-colonised preterm mice, sheep, rhesus macaques, baboons, and *Ureaplasma*-infected neonatal monocytes [4–6]. However, besides the apoptosis-modulating effects of acute *Ureaplasma* infection in pulmonary epithelial and endothelial cells [7, 8], further knowledge of the underlying mechanisms of *Ureaplasma*-driven preterm lung disease is missing.

The perinatal lung transition is critical in the neonate’s adaptation to air breathing. While intrapulmonary fluid is essential for fetal lung development, perinatally, this intrapulmonary fluid must be removed efficiently and quickly to fully establish postnatal lung function. This transition is impaired in preterm infants due to structural and functional lung immaturity [9]. The subsequent respiratory distress syndrome (RDS) is a leading cause of morbidity and mortality in preterm infants [10]. Besides surfactant deficiency, an immature epithelial Na^+^ channel (ENaC) expression has been acknowledged as a critical mechanism [9]. ENaC in the apical membrane compartment of alveolar type II (ATII) cells and the Na,K-ATPase in the basolateral membrane compartment promote the epithelial Na^+^ transport [11], which osmotically drives fluid absorption from the alveolar lumen into the interstitium and the circulation. Decreased alveolar fluid clearance in preterm infants [12] is most likely due to a lower ENaC expression [13]. Complete knockout of the α-ENaC subunit led to RDS and respiratory failure in newborn mice [14]. In ATII cells, ENaC comprises three homologous subunits, α-, β-, and γ-ENaC [11, 15], while the Na-K-ATPase contains α_1_- and β_1_-subunits [15].

The present investigation is the first to examine the influence of acute *Ureaplasma* infection on critical mechanisms of alveolar fluid clearance and perinatal lung transition of the immature lung, comprising epithelial Na^+^ transport, barrier integrity, metabolic activity, gene expression, and kinase signalling. We demonstrate a global inhibition of the epithelial Na^+^ transport in fetal alveolar cells upon *Ureaplasma* infection. *Ureaplasma*-driven phosphorylation of Erk1/2 most likely underlies this phenomenon. Moreover, the accumulation of *Ureaplasma*-derived ammonia (NH_3_) alone disrupts the Na^+^ transport. In line with this observation, flurofamide, an inhibitor of the *Ureaplasma*-specific urease enzyme, is identified as a counteragent preventing *Ureaplasma*-induced effects on epithelial Na^+^ transport. In summary, these data suggest a profound functional impairment of pulmonary epithelial cells upon *Ureaplasma* infection, potentially contributing to fluid accumulation and interfering with perinatal lung function in colonised preterm infants.

## MATERIALS AND METHODS

Detailed methods are described in the online supplement.

### Bacterial strains and culture conditions

*U. parvum* serovar 3 (Up3) and *U. urealyticum* serovar 8 (Uu8) were obtained from the American Tissue Culture Collection (ATCC, European distributor LGC Standards GmbH, Wesel, Germany; Up3: ATCC 27815, Uu8: ATCC 27618). Isolates were cultured in custom-made *in-house* medium as previously described [5, 6].

### Cell isolation and culture of FDLE cells and stimulation assays

FDLE cells, a model of preterm ATII cells, were isolated from rat fetal lungs as previously described [16]. Four days after isolation, cells were incubated with *Ureaplasma* spp., diluted 1:5 in cell culture medium for 3 or 24 hours. Moreover, FDLE were cultured with NaOH or NH_3_ at pH 8.0 for 24 hours. Sealing of the culture plates prevented the medium buffer from neutralising the pH value. Finally, co-incubation with flurofamide (10 µM) was used to inhibit the *Ureaplasma*-specific urease.

### Ussing chamber analyses

Ussing chamber measurements were performed as previously reported [17], following 3- and 24-hour incubation with *Ureaplasma*. Equivalent short-circuit currents (*I*_SC_) were determined every 20 sec by measuring transepithelial voltage (*V*_te_) and transepithelial resistance (*R*_te_) with a transepithelial current clamp (Physiologic Instruments, San Diego, CA) and calculating the quotient *I*_SC_ = *V*_te_/*R*_te_. After the *I*_SC_ reached a stable plateau (*I*_base_), amiloride (10 µM, # A7410, Sigma-Aldrich) was applied to the apical chamber. The amiloride-sensitive Δ*I*_SC_ (Δ*I*_amil_), a measure of ENaC activity, was calculated from the difference between *I*_base_ and the amiloride-insensitive *I*_SC_ (*I*_amil_). Maximal apical or basolateral Na^+^ permeability was determined by adding amphotericin B to the respective opposite compartment. At *I*_SC_ peak value, the amiloride-sensitive component (*amil*_max_) was determined by adding 10 µM amiloride to the apical compartment or ouabain (1 mM, # O3125, Merck) to the basolateral chamber to calculate the maximal ouabain-sensitive Na,K-ATPase activity (*ouab*_max_).

## RESULTS

### *Ureaplasma* infection inhibited Na^+^ transport in fetal alveolar cells

A 24-hour infection of FDLE cells with *U. urealyticum* (Uu8) reduced the epithelial Na^+^ transport by 70-90%. Ussing chamber analyses demonstrated that Na^+^ transport (*I*_base_) was significantly decreased by Uu8 infection compared with uninfected control cells (figure 1a). ENaC activity (Δ*I*_amil_) was also significantly reduced by Uu8-infection by more than 90%. In contrast, the epithelial barrier function assessed as *R*_te_ was unaffected by Uu8-infection. Furthermore, the amiloride insensitive current component, *I*_amil_, most likely Cl^−^ transport, was significantly reduced by Uu8 (figure 1a). To consider the capacities of ENaC and the Na,K-ATPase separately, the opposite membrane was permeabilised. The maximum amiloride-sensitive apical membrane permeability (*amil*_max_) was significantly decreased by Uu8 to approximately 35% (figure 1b). In addition, the maximum ouabain-sensitive Na,K-ATPase activity (*ouab*_max_) was significantly reduced upon Uu8 infection, although not to the same extent (figure 1c). In summary, Uu8 diminished the apical ENaC activity by more than 60% and the basolateral Na,K-ATPase activity by approximately 20% in FDLE cells upon 24-hour infection. In contrast, 3-hour incubation with Uu8 did not affect Na^+^ transport (figure 1d).

**Figure 1.**
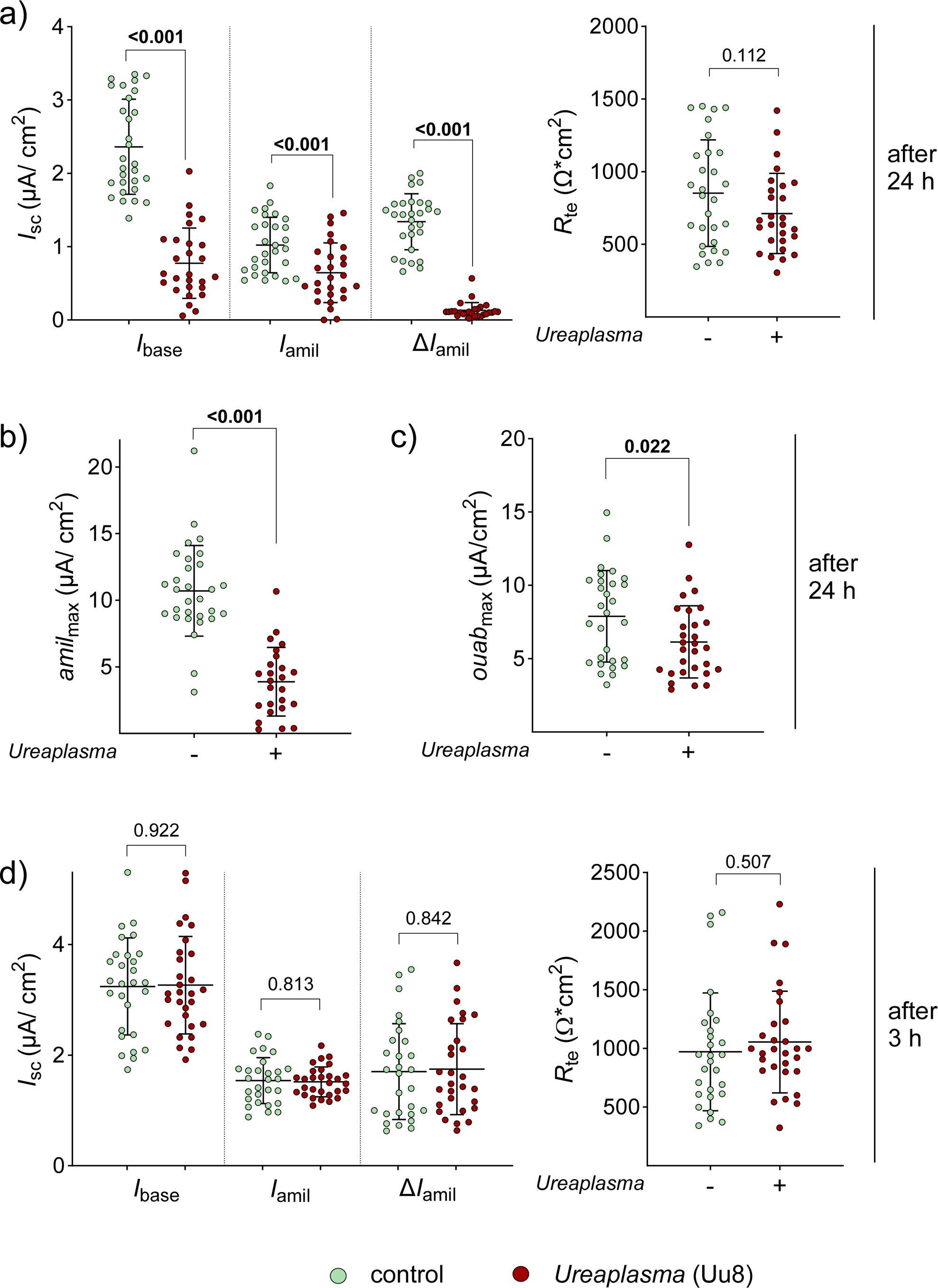
*Ureaplasma* infection strongly reduced epithelial Na^+^ transport in FDLE cells at 24-hour incubation. FDLE cells were infected with *U. urealyticum* (Uu8) for 24 **(a-c)** or 3 hours **(d)** before analyses. **a)** Uu8 infection significantly decreased Na^+^ transport and ENaC activity (*I*_base_, *I*_amil_, and Δ*I*_amil_). The barrier integrity (*R*_te_) of FDLE cells was not significantly altered (n=29) compared to uninfected controls (n=27). **b, c)** The maximal activity of each Na^+^ transporter was determined separately by permeabilising the opposite membrane. **b)** Uu8 infection significantly reduced the maximum apical Na^+^ permeability mediated by ENaC (*amil*_max_) (n=30) compared to controls (n=25). **c)** Moreover, maximum Na,K-ATPase activity (*ouab*_max_) was significantly reduced by *Ureaplasma* infection (n=29) compared to uninfected FDLE cells (n=30). **d)** Infection of FDLE cells with Uu8 for 3 hours did not affect *I*_base_, *I*_amil_, Δ*I*_amil_ or *R*_te_ (n=29) in comparison to control cells (n=28). The mean value for each group is represented by a horizontal line (±SD). Statistical analysis: unpaired t-test; *p*-value for each comparison is given in the graphs. Bold type represents statistical significance. *I*_base_, basal *I*_SC_; *I*_amil_, amiloride-insensitive *I*_SC_; Δ*I*_amil_, amiloride-sensitive *I*_SC_; *I*_SC_, short circuit current; *R*_te_, transepithelial resistance.

To determine if the effects observed for Uu8 were serovar-specific, measurements were repeated with Up3. As seen before, Up3 significantly decreased epithelial Na^+^ transport and ENaC activity in infected FDLE cells compared to uninfected controls (figure 2). Compared to Uu8, the reduction of Na^+^ transport by Up3 was less pronounced, attenuating ENaC activity by approximately 30%.

**Figure 2.**
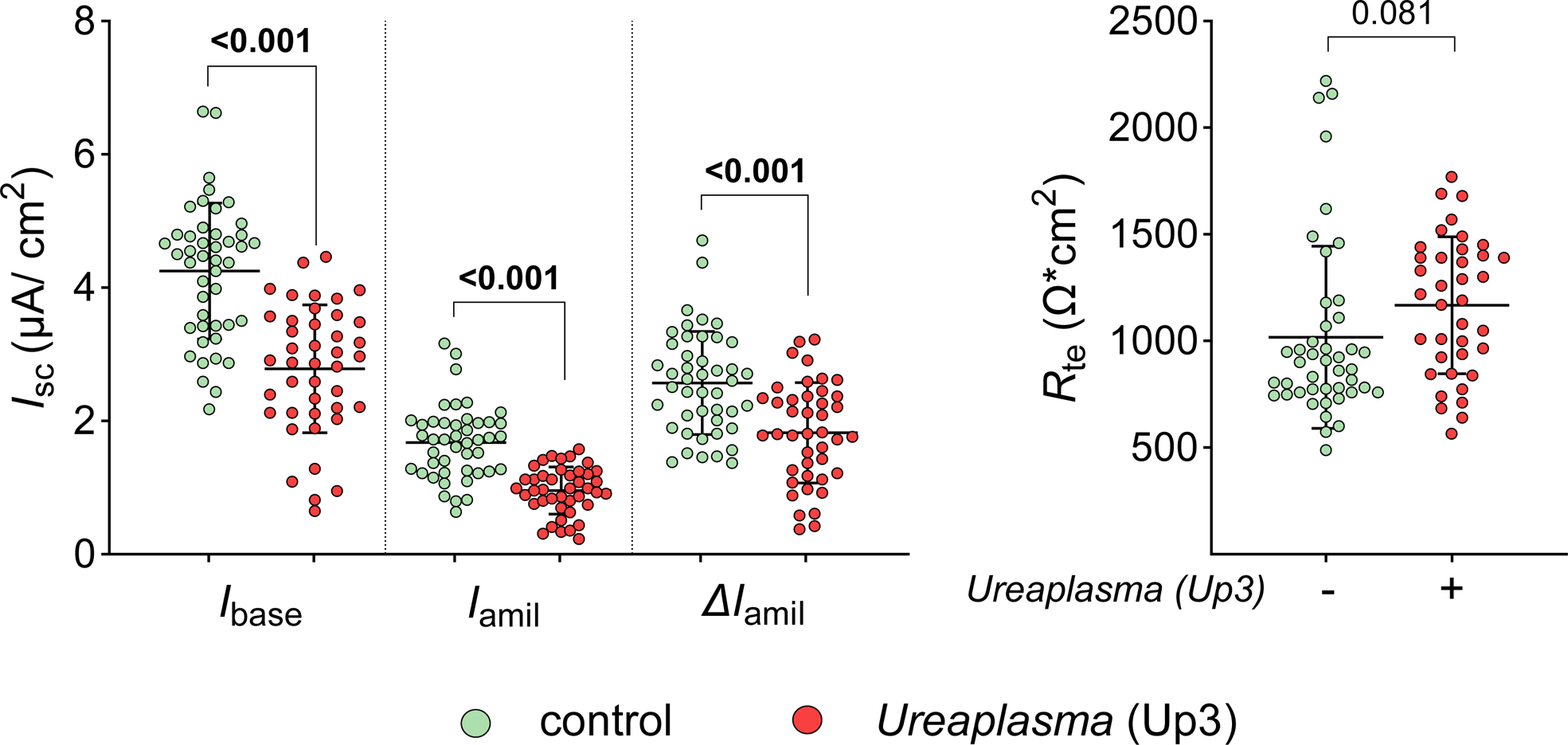
*U. parvum* (Up3) infection also strongly reduced epithelial Na^+^ transport in FDLE cells. Cells were infected with Up3 for 24 hours before analyses. In line with Uu8 infection, Up3 infection significantly decreased Na^+^ transport and ENaC activity (*I*_base_, *I*_amil_, and Δ*I*_amil_), while *R*_te_ was unaffected (Up3: n=45; control: n=42). The mean value for each group is represented by a horizontal line (±SD). Statistical analysis: unpaired t-test; *p*-value for each comparison is given in the graphs. Bold type represents statistical significance. *I*_base_, basal *I*_SC_; *I*_amil_, amiloride-insensitive *I*_SC_; Δ*I*_amil_, amiloride-sensitive *I*_SC_; *I*_SC_, short circuit current; *R*_te_, transepithelial resistance.

### *Ureaplasma* infection modulated gene expression and impaired metabolic activity in FDLE cells

Surprisingly, Uu8-infection significantly increased mRNA expression of the subunits *α-ENaC* by approximately 30% and *γ-ENaC* by almost 9-fold, while *β-ENaC* mRNA expression was unaffected (figure 3a). Moreover, Uu8-infection significantly increased mRNA expression of the *Na,K-ATPase-β_1_*-subunit by nearly 50%, while the *α_1_*-subunit level was not altered. In contrast, mRNA expression of the surfactant protein A (*Sftpa)* was decreased by approximately 70% and expression of *Sftpb* by around 30% (figure 3a). The opposite effect was observed for the *Sftpc* mRNA, which was increased by almost 40%. Taken together, Uu8-infection of FDLE cells increased the mRNA expression of several Na^+^ transporter subunits, in contrast to the observed functional impairment of the epithelial Na^+^ transport.

**Figure 3.**
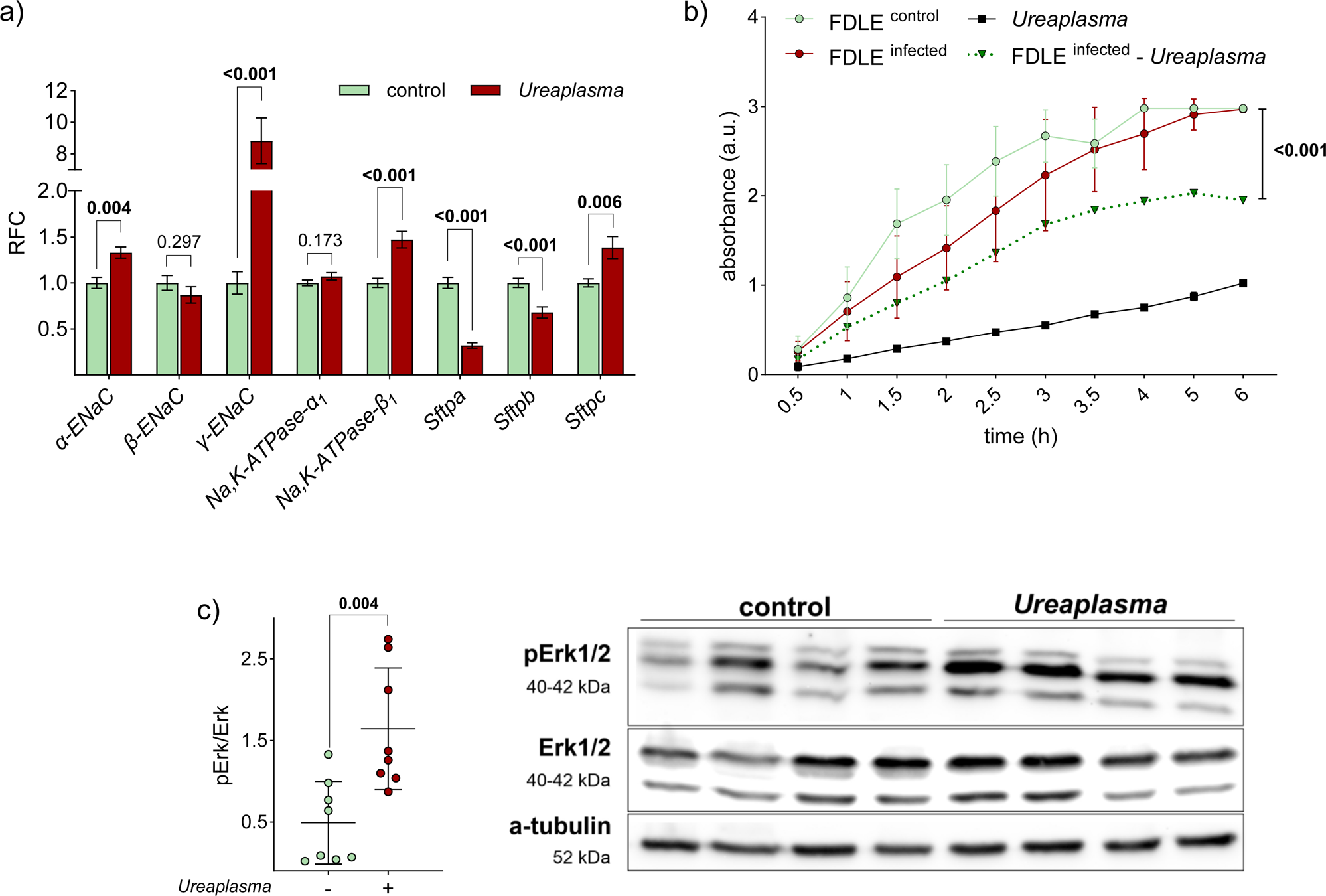
*Ureaplasma* infection significantly modulated mRNA expression of Na^+^ transporter subunits and surfactant proteins and affected the metabolic activity of FDLE cells and Erk1/2 signalling. **a, c)** FDLE cells were infected with Uu8 24 hours before analyses. **a)** Uu8 infection significantly increased mRNA expression of *α-* and *γ-ENaC*, *Na,K-ATPase-β_1_*, and *Sftpc*. In contrast, Uu8 infection downregulated *Sftpa* and *Sftpb* mRNA expression (n=12-18). Data are displayed as Mean (±SEM) of relative fold change (RFC). **b)** Metabolic activity was measured every 30 min to 1 hour in Uu8 infected FDLE cells (●), uninfected controls (●) and viable Uu8 isolates (▪). The dotted line (▾) represents the metabolic activity of Uu8-infected FDLE cells subtracted by the metabolic activity exhibited by Uu8 themselves. Based on this calculation, the metabolic activity was significantly lower in Uu8-infected FDLE cells than in uninfected controls (n=8-12 per group and time point). Data are displayed as mean (±SD). **c)** Uu8 infection significantly increased Erk1/2 phosphorylation (n=8) compared to controls (n=8). Representative Western blots give pErk1/2, total Erk1/2 and α-tubulin as loading control. The mean value for each group is represented by a horizontal line (±SD). Erk1/2, extracellular-signal-regulated kinases. Statistical analysis: **a, c)** unpaired t-test; **b)** repeated measures two-way ANOVA with Geisser-Greenhouse correction; p-value for each comparison is given in the graphs. Bold type represents statistical significance.

The metabolic activity of Uu8-infected FDLE cells was determined by WST-1 assay. Formazan products were measured in Uu8-infected FDLE cells, uninfected controls, and viable Uu8-isolates. No significant difference was observed in comparing Uu8-infected cells with uninfected controls (figure 3b). However, viable *Ureaplasma* isolates themselves reduced tetrazolium salts to formazan even in the absence of FDLE cells. We, therefore, subtracted the latter metabolic activity from results in Uu8-infected FDLE cells and found a significantly impaired metabolic activity in FDLE cells upon infection (figure 3b).

### *Ureaplasma* infection modulated phosphorylation of Erk1/2

To investigate whether key signalling pathways in FDLE cells were affected upon *Ureaplasma* infection, the phosphorylation of the signalling molecule Erk1/2 was analysed by Western blot. Erk1/2 is a well-known inhibitor of ENaC function. Uu8-infection strongly increased levels of pERK1/2 in FDLE cells by more than 3-fold (figure 3c), possibly contributing to the diminished ENaC activity.

### Ammonia mimicked *Ureaplasma*-driven effects on Na^+^ transport, gene expression and Erk1/2 phosphorylation

A 24-hour infection of FDLE cells with *Ureaplasma* caused a relevant pH shift (figure 4a). This was due to the metabolism of urea to ammonia (NH_3_) and the subsequent alkalisation of the culture medium. We tested whether the pH shift or NH_3_ itself was responsible for the observed *Ureaplasma*-driven effects. The pH shift to 8.0 using NaOH supplementation did not affect Na^+^ transport or ENaC activity, whereas adding NH_3_ to FDLE cells strongly reduced the Na^+^ transport (figure 4b). Taken together, *Ureaplasma*-induced ammonia and not the accompanying pH shift inhibited Na^+^ transport and ENaC activity. In contrast, the *Ureaplasma*-driven increase of α*-ENaC, γ-ENaC,* and *Sftpc* mRNA expression was mimicked by both the NaOH- and the NH_3_-induced pH shifts (figure 5a). Moreover, NaOH and NH_3_ both downregulated *Sftpa* mRNA expression. Finally, phosphorylation of Erk1/2 was increased by NaOH-mediated pH shift and NH_3_, but the effects of NH_3_ were significantly more pronounced (figure 5b).

**Figure 4.**
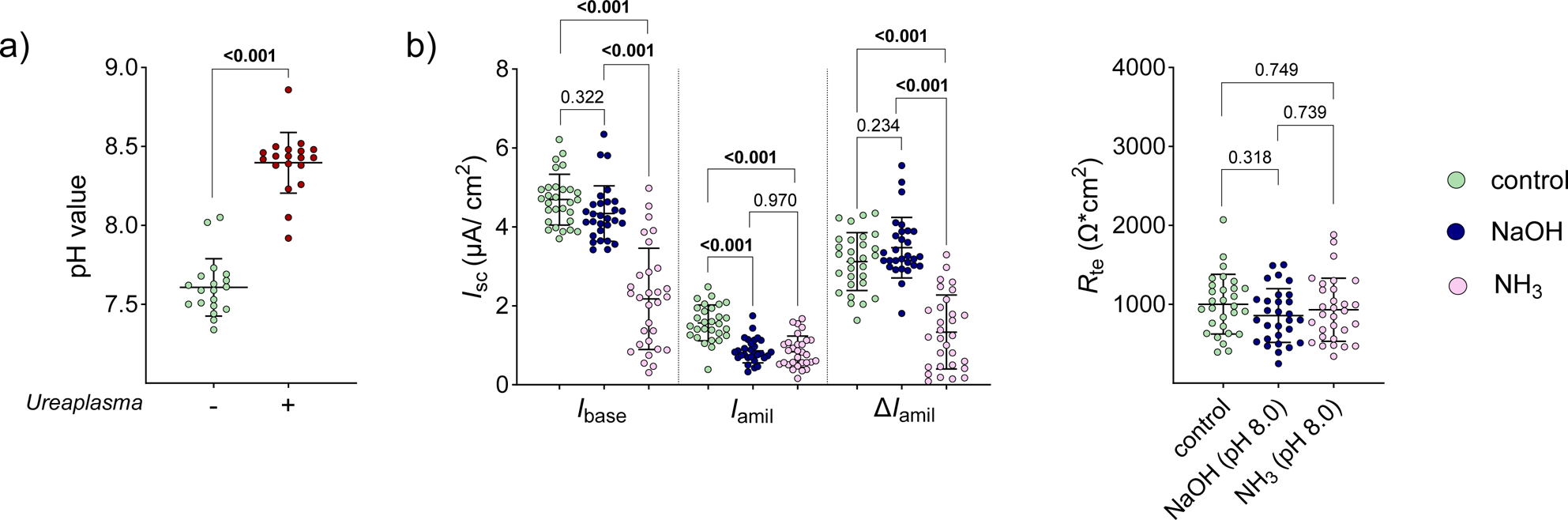
*Ureaplasma* infection shifted the pH value in infected FDLE cells due to ammonia (NH_3_) production, which in turn mimicked the effects of Uu8 infection to a large extent. **a)** Uu8 significantly increased the medium pH value in FDLE cells after 24 hours. **b)** The medium pH value of FDLE cells was shifted to pH 8.0 by adding either NaOH or NH_3_ and maintained for 24 hours. NH_3_ (n=30) significantly decreased Na^+^ transport and ENaC activity (*I*_base_, *I*_amil_, and Δ*I*_amil_) compared to control cells (n=29), while NaOH (n=29) only affected *I*_amil_. *R*_te_ was neither affected by NaOH nor NH_3_. The mean value for each group is represented by a horizontal line (±SD). Statistical analysis: **a)** unpaired t-test; **b)** one-way ANOVA with Tukey’s multiple comparison test; *p*-value for each comparison is given in the graphs. Bold type represents statistical significance. *I*_base_, basal *I*_SC_; *I*_amil_, amiloride-insensitive *I*_SC_; Δ*I*_amil_, amiloride-sensitive *I*_SC_; *I*_SC_, short circuit current; *R*_te_, transepithelial resistance.

**Figure 5.**
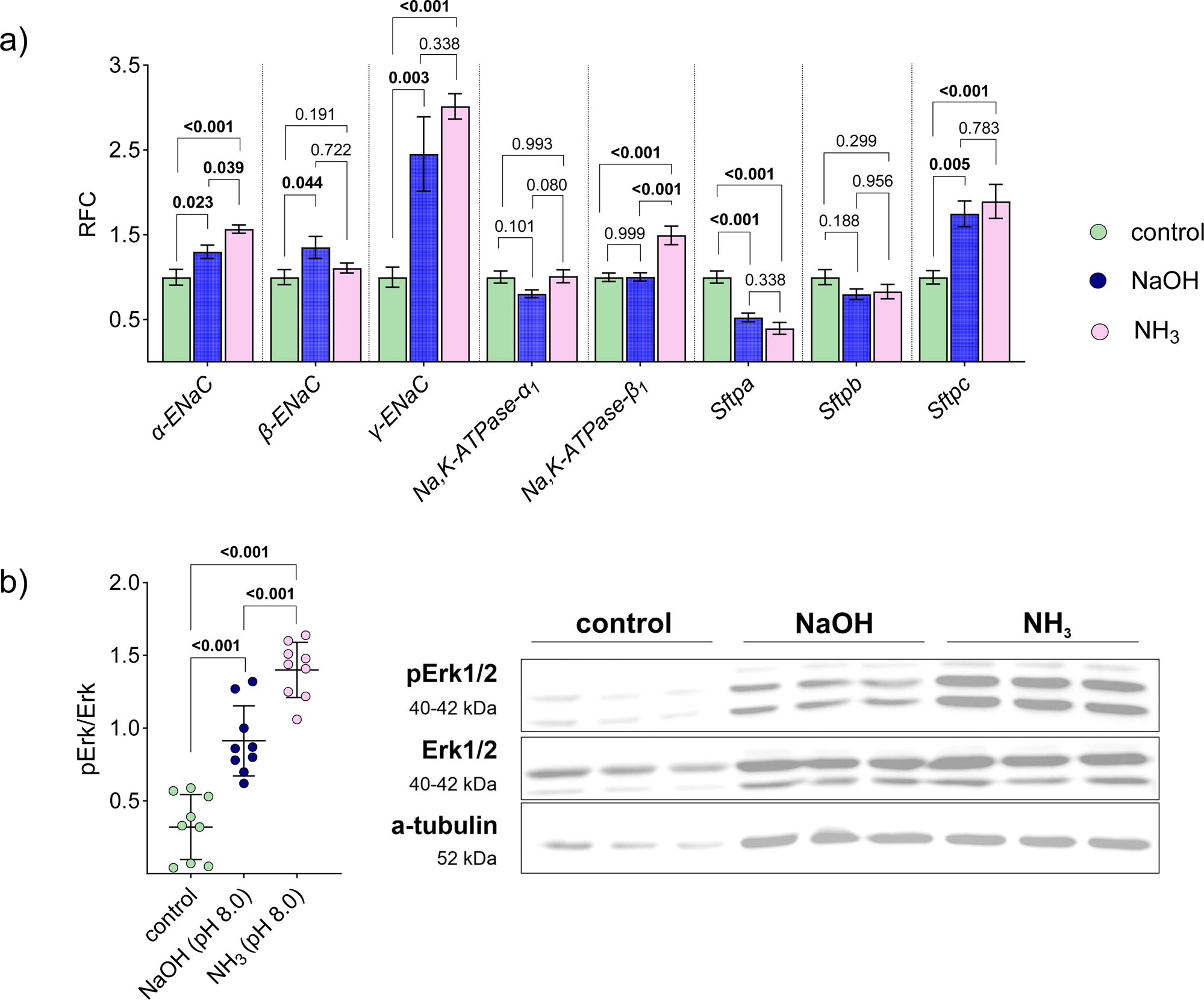
NaOH- and NH_3_-driven pH shift to 8.0 significantly modulated mRNA expression of Na^+^ transporter subunits and surfactant proteins and affected Erk1/2 signalling. FDLE cells were incubated with media containing NaOH and NH_3_ at pH 8.0 for 24 hours before analyses **a)** NaOH- and NH_3_-driven pH shift to 8.0 significantly increased mRNA expression of *α-* and *γ-ENaC*, *Na,K-ATPase-β_1_*, and *Sftpc*. In contrast, NaOH and NH_3_ downregulated *Sftpa* mRNA expression (n=11-12). Data are displayed as mean (±SEM) of relative fold change (RFC). **b)** NaOH- and NH_3_-driven pH shift to 8.0 increased Erk1/2 phosphorylation (n=9 each). Representative Western blots of pErk1/2, total Erk1/2 and α-tubulin are given. Erk1/2, extracellular-signal-regulated kinases. The mean value for each group is represented by a horizontal line (±SD). Statistical analysis: one-way ANOVA with Tukey’s multiple comparison test; *p*-value for each comparison is given in the graphs. Bold type represents statistical significance.

### The urease inhibitor flurofamide prevents *Ureaplasma*-driven impairment of Na^+^ transport and Erk1/2 phosphorylation

Flurofamide is a known inhibitor of bacterial urease and could, thus, interfere with the *Ureaplasma*-specific urease. According to our findings above, we hypothesised a role for urease as a key virulence factor in our experimental setting. To test the ability of flurofamide to restore Na^+^ transport in *Ureaplasma*-infected FDLE cells, cells were co-incubated with the urease inhibitor. *Ureaplasma*-driven inhibition of the Na^+^ transport was entirely prevented by flurofamide (figure 6a). Notably, flurofamide ameliorated ENaC activity in Uu8-infected cells to control levels. In addition, flurofamide attenuated the pH shift in Uu8-infected cells (figure 6b). The Uu8-induced increase of *α-ENaC*, *γ-ENaC* and *Sftpc* mRNA expression was significanly reduced by flurofamide (figure 7a). *Sftpa* mRNA expression was marginally elevated by flurofamide in Uu8-infected FDLE cells. Finally, flurofamide prevented the Uu8-induced phosphorylation of Erk1/2 (figure 7b), possibly contributing to the restored ENaC activity.

**Figure 6.**
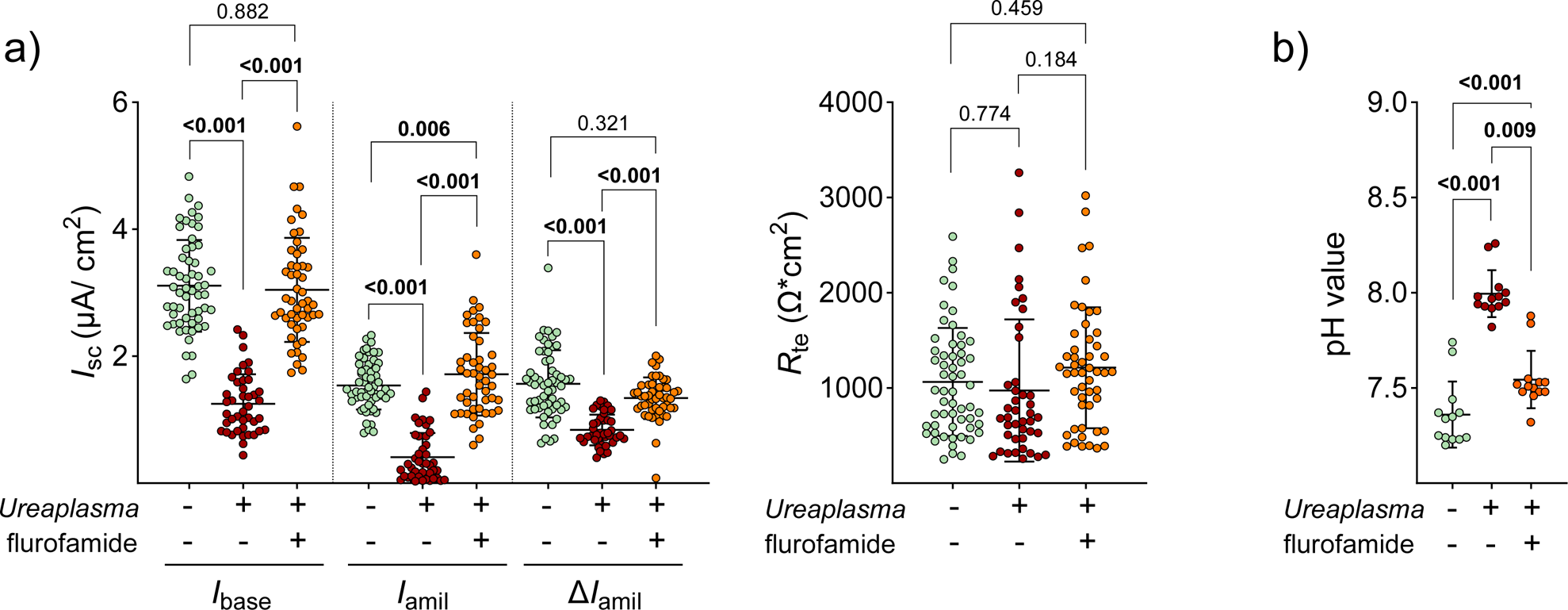
The presence of flurofamide restored the *Ureaplasma*-driven reduction of epithelial Na^+^ transport. FDLE cells were infected with Uu8 in the presence or absence of flurofamide (10 µM) for 24 hours. **a)** Uu8 infection (n=42) significantly decreased Na^+^ transport and ENaC activity (*I*_base_, *I*_amil_, and Δ*I*_amil_) compared to control cells (n=57). These findings were entirely restored by flurofamide (n=49). *R*_te_ was neither affected by Uu8 infection nor flurofamide. **b)** Flurofamide significantly downregulated media pH value in Uu8 infected FDLE cells (n=13 each). The mean value for each group is represented by a horizontal line (±SD). Statistical analysis: one-way ANOVA with Tukey’s multiple comparison test; *p*-value for each comparison is given in the graphs. Bold type represents statistical significance. *I*_base_, basal *I*_SC_; *I*_amil_, amiloride-insensitive *I*_SC_; Δ*I*_amil_, amiloride-sensitive *I*_SC_; *I*_SC_, short circuit current; *R*_te_, transepithelial resistance.

**Figure 7.**
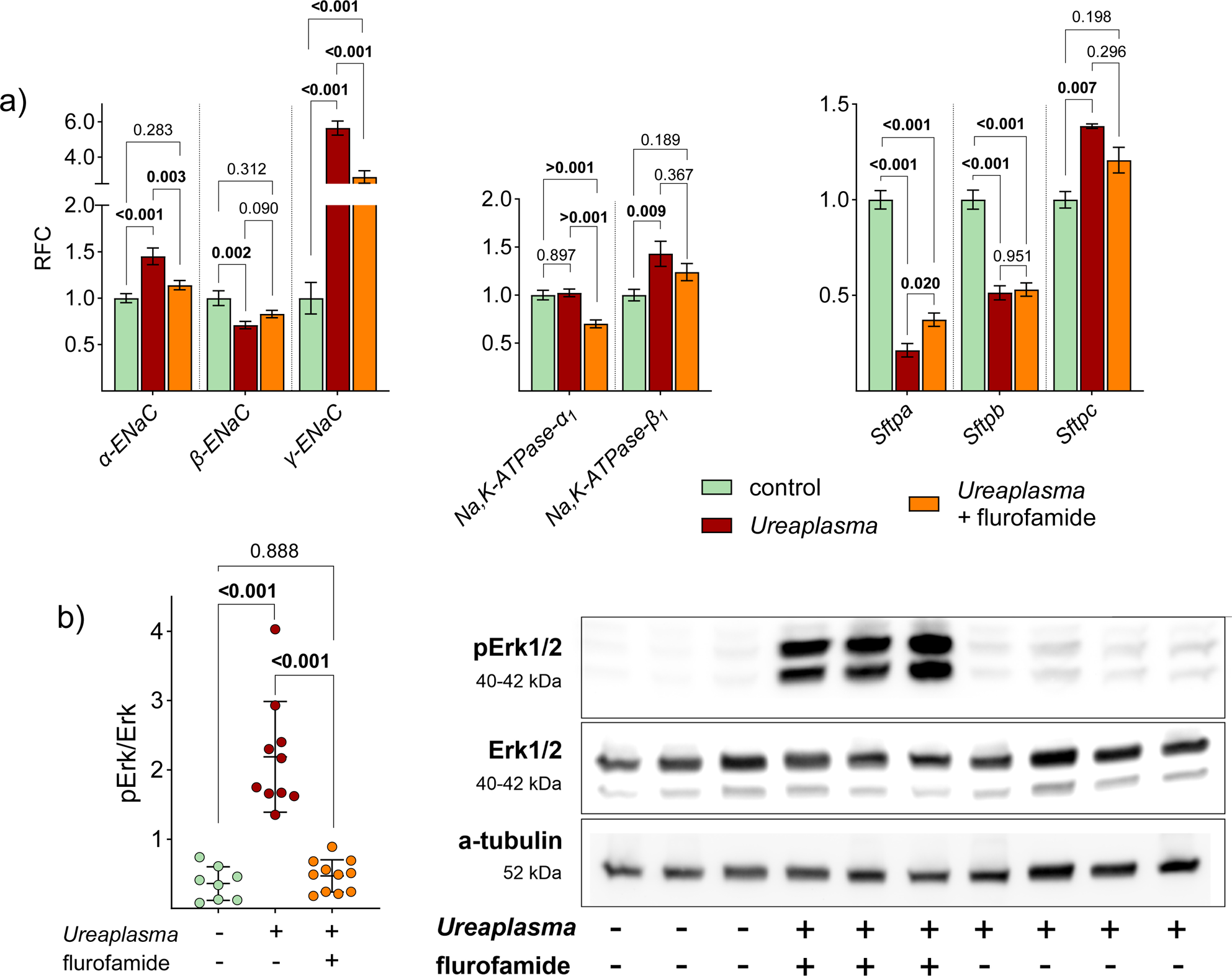
FDLE cells were infected with Uu8 in the presence or absence of flurofamide (10 µM) for 24 hours. **a)** Flurofamide had a marginal impact on Uu8-induced changes in mRNA expression of Na^+^ transporters and surfactant proteins: While Uu8 infection significantly increased mRNA expression of *α-* and *γ-ENaC*, *Na,K-ATPase-β_1_*, and *Sftpc*, flurofamide diminished the Uu8-induced increase of *α-* and *γ-ENaC*, and *Sftpc*. Uu8 infection downregulated mRNA expression of *Sftpa* and *Sftpb,* which was not affected by flurofamide (n=12 each). Data are displayed as mean (±SEM) of relative fold change (RFC). **b)** Addition of flurofamide prevented the Uu8-driven Erk1/2 phosphorylation. Uu8 infection significantly increased Erk1/2 phosphorylation, while phosphorylation was not seen in the presence of flurofamide (n=8 each). Representative Western blots of pErk1/2, total Erk1/2 and α-tubulin are given. Erk1/2, extracellular-signal-regulated kinases. The mean value for each group is represented by a horizontal line (±SD). Statistical analysis: one-way ANOVA with Tukey’s multiple comparison test; *p*-value for each comparison is given in the graphs. Bold type represents statistical significance.

## DISCUSSION

To our knowledge, this is the first study to describe a profound impairment of the epithelial Na^+^ transport in fetal alveolar cells upon acute *Ureaplasma* infection. Respiratory tract colonisation of preterm infants with *Ureaplasma* spp. has been frequently associated with adverse pulmonary short- and long-term outcome, in particular with the development of bronchopulmonary dysplasia [2, 3]. Since the epithelial Na^+^ transport is essentially involved in perinatal lung transition to air breathing, we sought to determine the potential impact of acute *Ureaplasma* infection on Na^+^ transport-associated mechanisms of alveolar fluid clearance.

In the present study, we could separate the effects on basolateral Na,K-ATPase, and apical Na^+^ channels by selective permeabilisation and observed a significant inhibition of the apical ENaC activity and a less profound reduction of the Na,K-ATPase activity induced by *Ureaplasma*. Notably, diminished epithelial Na^+^ transport in preterm infants [12] has been shown to contribute to the development of RDS [9]. According to our present findings, functional impairment of the Na^+^ transport in *Ureaplasma*-colonised preterm infants could further aggravate the critical respiratory situation for very preterm infants and worsen their clinical outcome.

Little is known about differences in virulence among certain *Ureaplasma* serovars [1, 18]. In the present study, we confirmed that both isolates, *U. parvum* and *U. urealyticum*, profoundly affected the epithelial Na^+^ transport. However, the *U. urealyticum* serovar exhibited a more potent inhibition, reducing the ENaC activity by 90%, while the *U. parvum* serovar achieved a lesser reduction of about 30%. The less profound effect seen for the *U. parvum* isolate might not correlate with a generally smaller effect of *U. parvum* on fetal alveolar cell function, but is most likely due to lower titers of Up3 in our experiments.

It is worth mentioning that the reduced activity of both Na^+^ transporters was not due to diminished mRNA expression. In fact, the mRNA expression of the *α-* and *γ-ENaC* subunits was increased, possibly reflecting a positive feedback mechanism. The functional relevance of each subunit might well explain the observed expression patterns of the ENaC subunits. In the fetal ovine lung, the *α-* and *γ-ENaC* subunit mRNA expression were shown to peak during labour [19], indicating that these ENaC subunits are critical at the developmental stage of the FDLE cells. On the other hand, overexpression of *β-ENaC* in mice decreases mucus clearance and reduces fluid in the postnatal lung, resulting in a cystic fibrosis-like phenotype [20]. The absent response of *β-ENaC* mRNA expression in FDLE cells in the present study might be explained by its relevance mainly for postnatal lung function.

This study showed an increased mRNA expression of the *Na,K-ATPase-β_1_* subunit in FDLE cells upon *Ureaplasma* infection. Overexpression of the *β_1_*-subunit was previously demonstrated to increase vectorial Na^+^ transport [16], with this subunit being the rate-limiting component in the assembly of the Na,K-ATPase [21]. In the present study, one may, thus, speculate that an upregulation of *Na,K-ATPase-β_1_* mRNA expression counteracts *Ureaplasma*-driven diminished Na^+^ transport.

The gene expression of surfactant proteins was modulated differently in *Ureaplasma*-infected FDLE cells. *Ureaplasma*-induced downregulation was observed for *Sftpa* and *Sftpb* mRNA expression, while *Sftpc* expression was increased. *Sftpa* contributes to the innate immune system by opsonisation processes, promoting the uptake of various microorganisms by phagocytic cell types [22]. Moreover, *Sftpa* protects the alveolar region from an uncontrolled inflammatory response by reducing proinflammatory Toll-like receptor signalling [22]. Subsequently, the observed *Ureaplasma*-induced reduction of *Sftpa* expression in our study could weaken alveolar defence mechanisms. This mechanism has not been described before. Of interest, *Ureaplasma*-driven downregulation of *Sftpa* might also impair the host’s immune defence against *Ureaplasma*. *Sftpa* was demonstrated to enhance ureaplasmacidal activity in murine alveolar macrophages [23]. *Sftpb* and *Sftpc* are essential for the surface-spreading properties of pulmonary surfactant. Still, only the lack of *Sftpb* is incompatible with life, while *Sftpc* is supposed to contribute to the full mechanical stabilisation of the alveolar structure [22]. In the present study, *Ureaplasma* infection downregulated the indispensable expression of *Sftpb,* potentially further contributing to preterm lung disease.

WST-1 assays demonstrated a significantly lower metabolic activity in *Ureaplasma*-infected cells that might indicate augmented cytotoxicity. These findings align with pro-apoptotic and apoptosis-modulating effects of acute *Ureaplasma* infection described in pulmonary epithelial and endothelial cells [7, 8]. In our WST-1 assays, we demonstrated that *Ureaplasma* isolates themselves reduced tetrazolium salts to formazan even in the absence of host cells. Subsequently, the *Ureaplasma*-inherent metabolic activity was subtracted from the results assessed in *Ureaplasma*-infected FDLE cells. Our findings align with genomic sequence analysis, suggesting that NADH, reducing the tetrazolium salt WST-1 in the given assay, contributes to several enzyme reactions in *Ureaplasma* metabolic pathways [24].

In our study, Western blot analyses revealed *Ureaplasma*-induced phosphorylation of Erk1/2, a signalling molecule significantly involved in regulating ENaC activity in previous studies. Acknowledging that phosphorylation of Erk1/2 has been shown to diminish ENaC activity in renal cells [25], we speculate that *Ureaplasma*-driven Erk1/2 phosphorylation is the underlying mechanism of the observed *Ureaplasma*-mediated inhibition of ENaC activity. In total, several regulatory mechanisms for ENaC activity have been described, including translocation from intracellular pools to the plasma membrane [26], preventing ENaC degradation by phosphorylation of the ubiquitin ligase NEDD4L (neural precursor cell expressed, developmentally downregulated 4) [27], and channel activation by direct phosphorylation [28]. Erk1/2 was suggested to directly phosphorylate the β- and γ-ENaC subunits, enhancing interaction with NEDD4L and subsequent channel degradation [29]. Due to a lack of suitable antibodies against rat ENaC, we did not perform analyses of Na^+^ transporter protein expression, which is a limitation of this study. Therefore, we cannot exclude that (post)translational effects might also contribute to the downregulated Na^+^ transport in *Ureaplasma*-infected cells.

One potential virulence factor of *Ureaplasma* often discussed in the context of *Ureaplasma*-host cell interaction is the urease-mediated production of ammonia and the accompanying pH shift [1, 3]. Although frequently considered, this potential virulence factor has not been comprehensively investigated. In the present study, we determined whether the *Ureaplasma*-derived NH_3_ or the NH_3_-driven pH shift disrupted the epithelial Na^+^ transport. While a NaOH-mediated pH shift did not affect ENaC activity, NH_3_ inhibited the epithelial Na^+^ transport in FDLE cells, mimicking *Ureaplasma*-driven effects. Moreover, NH_3_ affected mRNA expression of Na^+^ transporters and surfactant proteins and induced Erk1/2 phosphorylation similar to *Ureaplasma* isolates. Therefore, it can be assumed that *Ureaplasma*-driven NH_3_ represents a key virulence factor in *Ureaplasma*-alveoli interaction impairing alveolar epithelial function. To further delineate the relationship between Na^+^ transport and *Ureaplasma*-induced NH_3_ production, we inhibited the bacterial urease with flurofamide. Following our hypothesis, flurofamide completely restored Na^+^ transport and ENaC activity in *Ureaplasma*-infected FDLE cells. In addition, flurofamide prevented the *Ureaplasma*-driven phosphorylation of Erk1/2. This aligns with studies confirming that flurofamide inhibits ammonia production from urea by intestinal microorganisms *in vitro* [30]. Notably, *Ureaplasma*-driven ammonia has been implicated in the pathogenesis of hyperammonemia syndrome in infected immunocompromised patients [31, 32], leading to neurological dysfunction and frequent mortality [33, 34]. A mouse model confirmed fatal *Ureaplasma-*induced hyperammonemia in the context of immunosuppression [35]. Flurofamide reduced blood ammonia levels in these *Ureaplasma*-infected hyperammonemic mice [36]. In line with these data, our findings of complete prevention of *Ureaplasma*-driven effects by flurofamide make this substance or other urease-inhibiting agents promising candidates for future non-antibiotic treatment strategies in *Ureaplasma*-colonised preterm infants. This is especially important since no treatment is available for improving Na^+^ transport and alveolar fluid clearance at the time of perinatal lung transition.

Taken together, our study describes the *Ureaplasma*-driven inhibition of Na^+^ transport and ENaC activity in fetal alveolar cells, most likely due to altered Erk1/2 signalling. *Ureaplasma*-induced production of NH_3_ by urea degradation was identified as a key underlying mechanism. NH_3_ addition to FDLE cells mimicked most *Ureaplasma-*driven effects on FDLE cell function. Moreover, flurofamide-mediated prevention of *Ureaplasma*-driven NH_3_ production completely restored Na^+^ transport and ENaC activity, as well as Erk1/2 activity, confirming a contributing role of the *Ureaplasma*-specific urease. To the best of our knowledge, this is the first study to show a functional impairment of pulmonary epithelial cells upon *Ureaplasma* infection and to reveal potential mechanisms of *Ureaplasma*-driven lung disease in preterm infants. According to our findings, perinatal inhibition of the epithelial Na^+^ transport in *Ureaplasma*-colonised infants would contribute to significant lung fluid accumulation – and would further impair lung transition and function, especially in the most immature infants. Ultimately, these data underline the relevance of *Ureaplasma* spp. as true pathogens in preterm infants, and they shed light on the role of urease-inhibiting drugs, such as flurofamide, as potential therapeutic strategies in colonised preterm infants early in life.

## Acknowledgements

We thank Jessica Löffler, Sören Pietsch, and Chiara Becattini for their excellent technical assistance and Nadine Dietze for providing and organising the microbiological laboratory facilities. Parts of the doctoral theses of C.B. Rieger and E. Paluszkiewicz are incorporated into this manuscript.

## Ethics approval

All experimental protocols were approved by the institutional review board (IRB: Landesdirektion Leipzig, Leipzig, Germany, permit number: TVV15/22).

## Author contributions

Conceptualisation: M. Laube and K. Glaser. Methodology: M. Laube and K. Glaser. Formal analysis: M. Laube. Investigation: C.B. Rieger and E. Paluszkiewicz Data curation: C.B. Rieger and E. Paluszkiewicz. Writing (original draft): M. Laube and K. Glaser. Visualisation: M. Laube. Supervision: M. Laube, U.H. Thome and K. Glaser. Project administration: M. Laube and K. Glaser. Funding acquisition: M. Laube and K. Glaser.

## Data availability statement

All data generated or analysed in this study are either included in this article, uploaded as online supplemental information or can be obtained from the corresponding author on reasonable request.

## Supplement

### *Ureaplasma* culture conditions

Frozen stocks were prepared from mid-logarithmic-phase broth culture and stored at −80◦C until use. For each experiment, both isolates were inoculated 1:10 with in-house medium (“broth”) containing 82% autoclaved PPLO medium (Becton, Dickinson & Company, Franklin Lakes, NJ, USA), 11% heat-inactivated horse serum (v/v), urea (186 mM, Merck, Taufkirchen, Germany) and 1% phenol red (Merck), filtered through a 0.2-micron membrane and adjusted to pH 6.5. Ten-fold serial dilutions of both strains were incubated overnight to obtain titers of 1×10^7^ to 1×10^9^ colour-changing units (CCU)/mL of viable Up3 and Uu8. BacTiter-Glo^TM^ Microbial Cell Viability Assays (Promega GmbH, Walldorf, Germany) were used to screen for ATP production from viable organisms. Determination of CCUs was performed in 96 well plates (Greiner, Frickenhausen, Germany) as described earlier [1], paralleling each experiment. Corresponding DNA copy numbers of Uu8 and Up3 DNA were determined by quantitative PCR using the following primers Uu8-s 5’ CTGACCACGTAGTGGAAGGG 3’ and Uu8-as 5’ CAACTTGGATAGGACGGTCACC 3’ as well as Up3-s 5’ GCTGACCTAATGCAATCTGCTCG 3’ and Up3-as 5’ GGACTAAATTGACTTGATGATCCTG 3’.

### Cell isolation and culture of FDLE cells

All experimental protocols were approved by the institutional review board (IRB: Landesdirektion Leipzig, Leipzig, Germany, permit number: TVV15/22). All methods were performed following relevant guidelines and regulations. Sprague-Dawley rats were bred at the Medical Research Center of the University of Leipzig. Rats were housed in a room with a 12-hour light/dark cycle at constant temperature (22 °C) and humidity (55%). Food and water were provided ad libitum. Pregnant rats were euthanised by pentobarbital injection on gestation days E20-21 (term E=22).

After mechanical dissociation of fetal lungs, the cell suspension was enzymatically digested with 0.125% trypsin (Life Technologies, Darmstadt, Germany) and 0.4 mg/mL DNAse (CellSystems, Troisdorf, Germany) in HBSS (Life Technologies) for 10 min at 37°C. Then, the cells were incubated with MEM containing 0.1% collagenase (CellSystems) and DNAse at 37°C for another 15 min. To remove contaminating fibroblasts, cells were plated twice for one hour at 37°C in cell culture flasks. For electrophysiological measurements, FDLE cells were seeded onto permeable Snapwell inserts (surface area 1.1 cm^2^, Costar 3407; Corning, Corning, NY, USA) at a density of 10^6^ cells per insert. For RNA isolation, cells were seeded onto larger inserts (ThinCert, #657641, 4.6 cm^2^ surface area, Greiner Bio-One, Frickenhausen, Germany) at a density of 2 x 10^6^ cells per insert. Standard cell culture medium consisted of MEM with 10 % fetal bovine serum (FBS, Biochrom, Berlin, Germany), glutamine (2 mM, Life Technologies), and antibiotic-antimycotic (Life Technologies). The cell culture medium was changed daily.

### Ussing chamber analyses

Only monolayers with a transepithelial resistance (*R*_te_) exceeding 300 Ω·cm^2^ were included in the analyses. Electrophysiological solutions consisted of: 145 mM Na^+^, 5 mM K^+^, 1.2 mM Ca^2+^, 1.2 mM Mg^2+^, 125 mM Cl^−^, 25 mM HCO^3−^, 3.3 mM H_2_PO ^−^ and 0.8 mM HPO ^2−^ (pH 7.4). While 10 mM glucose was used for the basolateral solution, 10 mM mannitol was used in the apical solution. During measurements, the solutions were continuously gassed with carbogen (5% CO2 and 95% O2). For permeabilised membrane measurements, the *I*_SC_ was measured every 5 sec with a transepithelial voltage clamp. The apical Na^+^ permeability was determined by adding amphotericin B (a pore-forming antibiotic, 100 µM, # A-4888, Merck) to the basolateral compartment after the *I*_SC_ reached a plateau (*I*_base_). For this experimental setup, 140 mM of basolateral Na^+^ was replaced by 116 mM N-methyl-D-glucamine (NMDG^+^, # M-2004, Merck) and 24 mM choline, generating a 145:5 apical-to-basolateral Na^+^ gradient. Following amphotericin B addition and *I*_SC_ peak, the amiloride-sensitive component (*amil*_max_) was determined by adding 10 µM amiloride to the apical compartment. Permeabilising the apical membrane with amphotericin B (10 µM) loads the cell interior and the Na,K-ATPase with Na^+^, which enables the determination of the maximal ouabain-sensitive Na,K-ATPase activity (*ouab*_max_). Amiloride and ouabain stock solutions were prepared in water.

### mRNA expression analyses

RNA isolation was done following a 24-hour incubation with *Ureaplasma* spp. using the Purelink RNA Mini Kit (Life Technologies) and DNAse I (Life Technologies) according to the manufacturer’s instructions. For reverse transcription, 1 g of RNA was pre-annealed with Oligo(dT)18 primers (Fisher Scientific GmbH, Schwerte, Germany), followed by the addition of Superscript III (Life Technologies) for 1 hour at 55 °C and 15 min at 75 °C. The Express Greener QPCR Uni Kit (Fisher Scientific GmbH), gene-specific primers (table 1) [2], and the CFX 96 Real-Time system (Bio-Rad, Munich, Germany) were used to perform real-time quantitative PCR (RT-qPCR). Target-specific plasmid DNA was used as the internal standard for absolute quantification. The resultant molecular concentrations were normalised to the mitochondrial ribosomal protein S18a (Mrps18a) reference gene. Mrps18a’s consistent expression was confirmed using other widely used reference genes. The relative standard curve approach was used to calculate the mRNA levels, expressed as the respective control’s relative fold change (RFC). To control the specificity of the PCR reaction, melting curves and gel electrophoresis of PCR products were frequently carried out.

**Table 1.**
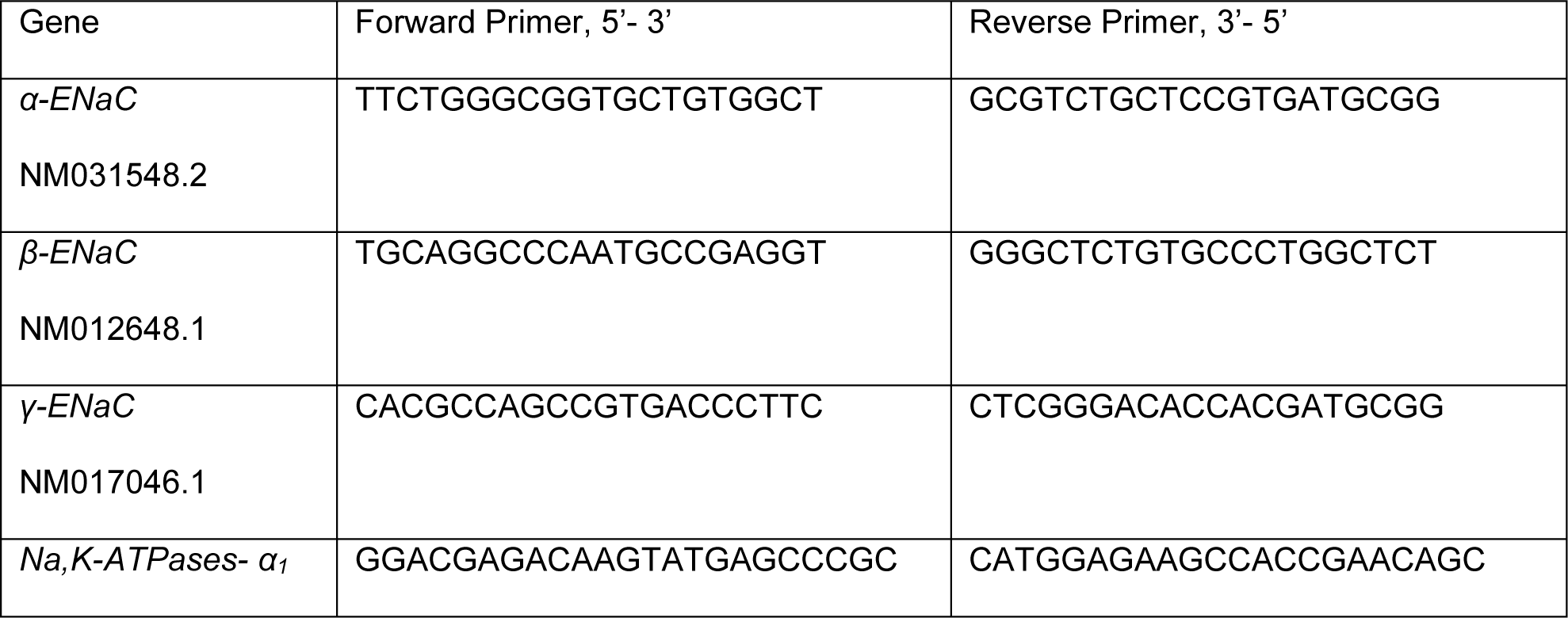
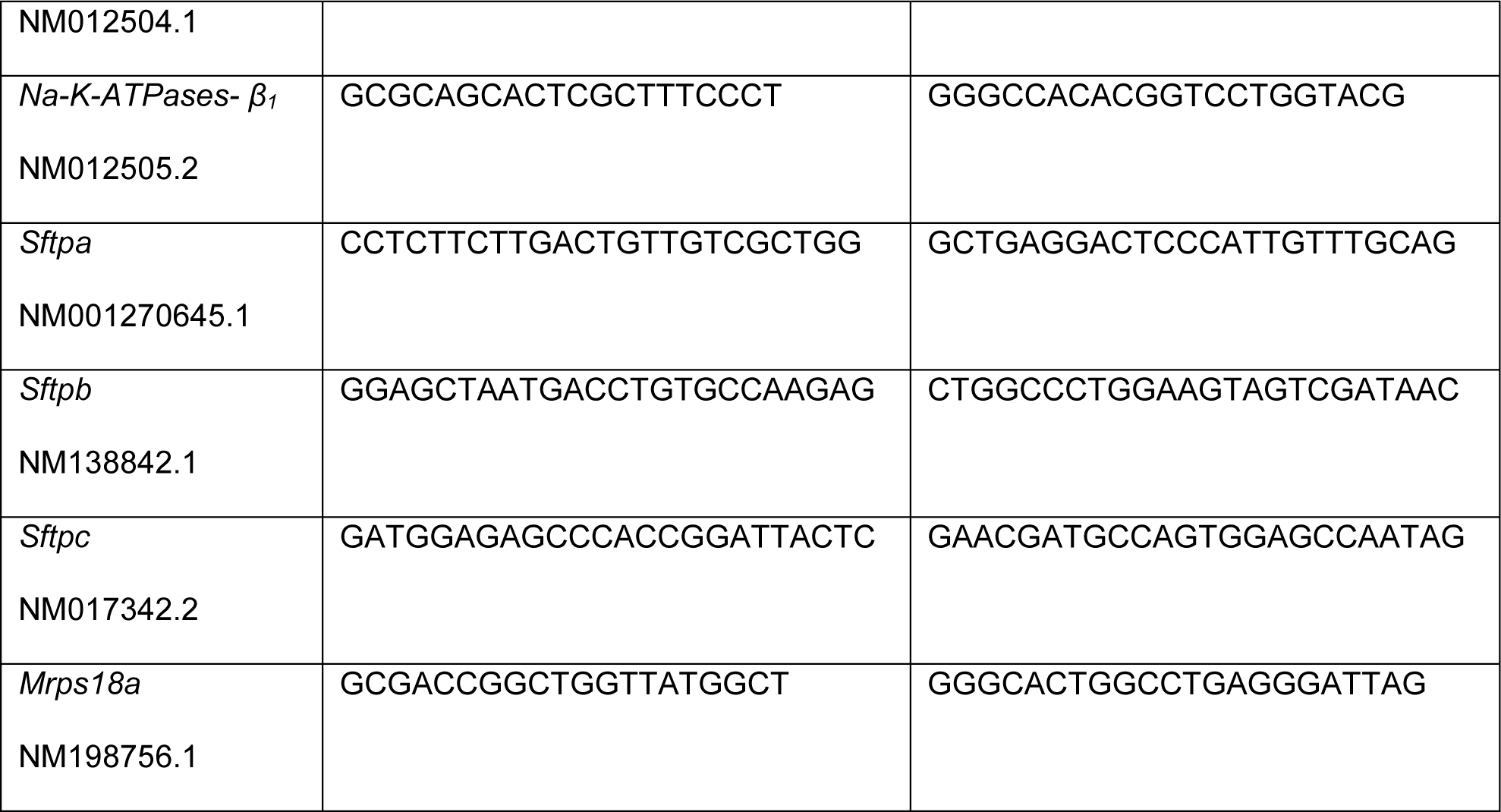
Primer sequences.

### WST-1 assay for cell proliferation and viability

Nicotinamide adenine dinucleotide (NAD) is involved in redox reactions, and NADPH reduces the tetrazolium salt WST-1 [(4-[3-(4-Iodophenyl)-2-(4-nitro-phenyl)-2H-5-tetrazolio]-1,3-benzene sulfonate] to formazan, which is measured spectrophotometrically in WST-1 assays (Roche, Mannheim, Germany). Metabolic activity comprising cell proliferation and viability was determined in *Ureaplasma*-infected FDLE cells, uninfected controls and cultured, viable *Ureaplasma* isolates. FDLE cells were seeded in 96 well plates at a density of 2 x 10^4^ cells per well and infected with *Ureaplasma* (Uu8). After 24 hours, the medium was changed, and WST-1 reagent (Roche, Mannheim, Germany) was added according to the manufacturer’s instructions. Absorbance at 450 nm was measured regularly every 0.5 to 1 hour.

### Western blot analyses

FDLE cell inserts were placed on ice, and protein lysates were prepared and analysed as described elsewhere [2]. Phosphorylation of Erk1/2 (extracellular-signal-regulated kinases) was studied using antibodies against phospho-Erk1/2 at Tyr202/Tyr204 (# 9101, Cell Signaling Technology), and Erk1/2 (# 9102, Cell Signaling Technology). Furthermore, the expression of α-Tubulin (11H10, # 2125, Cell Signaling Technology) was used as a reference. Suitable secondary antibodies conjugated to horseradish peroxidase (HRP) were used to detect primary antibodies. HRP activity was analysed by enhanced chemiluminescence (ECL, Amersham, Piscataway, NJ, USA) on x-ray film, and band intensity was measured by densitometry using Image-J (National Institutes of Health (NIH), Bethesda, MD, USA).

### Statistical analyses

Statistical analyses were performed using GraphPad Prism software (GraphPad Prism 9.1.1, https://www.graphpad.com, GraphPad Software, La Jolla, CA, USA). Data are expressed as mean ± standard deviation (SD) or standard error (SEM). Unpaired t-test, one-way ANOVA followed by Turkey’s multiple comparison post-hoc test, and two-way ANOVA with Geisser-Greenhouse correction were performed comparing expression levels among study groups. Statistical significance was defined as *p*-value < 0.05.

